# Family matters! The influence of family, peers, mentors, and professors on STEM college students’ motivational beliefs and career decision making

**DOI:** 10.1101/2025.01.22.633242

**Authors:** Trevor Tuma

## Abstract

Support from family, peers, mentors, and professors has a crucial role in the development of students’ motivational beliefs about learning. However, the *relative* importance of each of these social supports in affecting students’ motivation, and the precise processes through which these social factors motivate students, are not well understood at the college level. The present study examined how family, peers, mentors, and professors shaped science, technology, engineering, and mathematics (STEM) college students’ motivational beliefs for career decision making, defining motivational beliefs in accordance with situated expectancy-value theory. We collected survey and open-ended responses from a large sample of college students (*n* = 2,229) in 42 unique STEM courses at a large, public southeastern university in the United States. We examined how, why, and for whom different social supports influenced students’ task-value and/or competence related motivational beliefs for career-decision making. Results indicated that students most often selected interactions with family members as influencing their career decision-making, both in terms of overall influences and as the strongest influence, relative to interactions with peers, mentors, or professors. For all social influences, most students reported being attracted in a positive way towards certain careers as a result of social interactions, as opposed to socializers making students disenchanted with careers. Most students described how social supports influenced motivational beliefs related to task values, with comparatively fewer referencing social influences on competence-related beliefs. Findings highlight the important role of multiple social supports, particularly family, in shaping students’ motivational beliefs and career decision-making throughout emerging adulthood.

**Educational Impact and Implications Statement:** Preparing a qualified and diverse STEM workforce requires a robust understanding of the factors that influence college students’ motivational beliefs and decisions about their career paths. We examined how, why, and for whom support from family, peers, mentors, and professors impacted STEM college students career motivation and decision making, using both qualitative and quantitative data. Results suggest that family plays a particularly crucial role in shaping college students’ career motivation, especially by helping them see the positive value of particular career paths. Findings help illustrate the importance of different kinds of social support in guiding career choices throughout emerging adulthood.

## Introduction

Attracting and retaining students in science, technology, engineering, and mathematics (STEM) fields remains an urgent national priority (Brewer & Smith, 2011; Olson & Riordan, 2012). For example, in the United States, more than half of the undergraduate students who intend to major in a STEM discipline will not persist to graduate with a STEM degree (Chen, 2013; Graham et al., 2013). Such trends limit the pool of individuals who have the educational experiences, skills, and knowledge necessary to pursue a career in STEM. Furthermore, students from racially and ethnically excluded backgrounds (e.g., Alaskan Native, American Indian, African American, Hawaiian/Pacific Islander, Hispanic or Latino/Latina) leave STEM fields at higher rates than students from well-represented backgrounds (e.g., Asian, White) (Estrada et al., 2016). Even students who remain in STEM fields often end up shifting their career plans during college (Seymour et al., 2019). About half of students who remain in STEM career paths across college will change their career plans *within* STEM (i.e., changing from wanting to be a doctor to a social worker) (Authors, 2021a; Authors, 2024a). College students who change their career plans within STEM in turn report lower motivation, satisfaction, and certainty of persisting with their new career plans compared to those with consistent career plans (Authors, 2024a).

In order to support STEM workforce development, it is essential to understand why students who start college interested in STEM career paths may change their plans to pursue other careers both inside and out of STEM. From a developmental perspective, the college years are especially critical for supporting STEM workforce development, because individuals make impactful career-related decisions (e.g., majors, internships, graduate programs) during this time. In the present study, we examined one key aspect of career plan development in college: the role of socializers (i.e., family, peers, mentors, and professors) in shaping college students’ STEM career motivation. Our goal was to build on prior work in this area by leveraging both quantitative and qualitative data from a large sample of college students. We examined which socializers college students reported to be most impactful in shaping career planning, which motivational beliefs students thought socializers affected most readily, how students thought different socializers shaped their motivation, and whether trends differed for students of different genders, racial/ethnic backgrounds, or years in school.

### Situated Expectancy-Value Theory and Career Decision-Making

Mounting evidence suggests that students’ motivational beliefs are powerful predictors of their persistence and career decision-making in STEM fields (Cromley et al., 2016; Lazarides & Lauermann, 2019; Wang & Degol, 2013). Eccles and colleagues’ expectancy-value theory (Eccles, 1983), recently renamed situated expectancy-value theory (SEVT) (Eccles & Wigfield, 2020), continues to be one of the most prominent theories scholars have used as a framework for understanding students’ motivational beliefs for academic decision-making. SEVT posits that individuals’ motivation to engage in certain tasks and subsequent outcomes, such as deciding to pursue a STEM career, are influenced by their expectancies to succeed at a task (e.g., Can I do this?) and subjective perceptions of values relating to the task (e.g., Do I want to do this?). When students have higher expectancies and values for a particular task, they are more likely to engage in that task and perform well (Eccles & Wigfield, 2020). An individual’s expectancies and task values are thought to be contextualized or situated within particular contexts, suggesting that they manifest uniquely, dynamically, and synergistically within different learning environments and in response to different social and contextual experiences (Eccles & Wigfield, 2020).

A considerable volume of research has shown that students’ expectancies (or closely related competence-related beliefs such as academic self-concept and self-efficacy) and the different aspects of subjective task values predict their academic achievement and choices of subsequent activities such as course-taking or college majors (Authors, 2021b; Canning et al., 2018; Gaspard et al., 2019; Musu-Gillette et al., 2015; Simpkins et al., 2006). Students’ competence-related beliefs and task values are both key predictors of students’ intentions to pursue a STEM major or actual decisions to pursue a STEM-related career path (Guo et al., 2015; Lauermann et al., 2017; Musu-Gillette et al., 2015; Perez et al., 2014; 2019). This large body of research together illustrates that SEVT is a powerful and useful theoretical framework for understanding the key motivational beliefs that shape students’ career development and decision making.

Situated expectancy-value theory emphasizes the important role of socialization processes in shaping individuals’ motivational beliefs (Eccles & Wigfield, 2020; Anderman & Kaplan, 2008). Theoretically, individuals’ motivational beliefs for learning are thought to be affected by both broad social environments in which students learn, such as family, educational, and cultural contexts, and by socializer’s beliefs and behaviors that are directly communicated to students (Eccles, 2007; Jodl et al., 2001). These socialization processes can influence students in multiple directions, with students possibly becoming attracted to a particular career in a positive way after experiences with a socializer or becoming disenchanted away from a particular career as a result of social influences (Authors, 2021b). Sometimes socializers may enhance individuals’ beliefs in their competence and/or increase an individuals’ perceived value of pursuing a certain task, thus attracting them towards a particular activity or career path (e.g., if a student’s friend tells the student that they can succeed at a career path, the friend may make the student feel more confident). However, social influences also can *reduce* competence-related beliefs and/or task values and make the student disenchanted with a career opportunity (e.g., if a child grows up in a familial environment where a particular career path is not perceived as worthwhile, this may reduce the student’s task value perceptions for that path).

### Major Socializers of Students’ STEM-Related Motivation

The idea that students’ beliefs are shaped through social influences is not unique to SEVT, with a very large body of research inside and outside the field of educational psychology illustrating the powerful role of social influences on individuals’ beliefs, feelings, and behavior (Cialdini & Trost, 1998; Mason et al., 2007; Nolan et al., 2008; Schultz et al., 2007; Wentzel & Asher, 1995). This literature has shown that many types of social supports shape students’ motivation to choose particular career paths throughout childhood, adolescence, and emerging adulthood. Family members, teachers, and peers are arguably considered to be the most impactful social agents for shaping students’ motivational beliefs and subsequent achievement-related choices (Eccles et al., 2013; Wentzel, 1998). We discuss each socializer briefly in turn.

#### Family members

The nature and quality of students’ relationships with family members (particularly parents, but also siblings and other relatives) can have a profound impact on their educational experiences and academic achievement (Barger et al., 2019; Collins & Steinberg, 2008; Fan & Chen, 2001; Hill & Taylor, 2004; Jeynes, 2003, 2005; Pomerantz et al., 2007, 2012; Steinberg & Morris, 2001). Research has indicated specifically that parental involvement influences children’s achievement by shaping children’s motivational beliefs (Grolnick et al., 1991; Grolnick & Slowiaczek, 1994). Other research has shown that interactions between family members and children can predict students’ motivational beliefs (Cheung & Pomerantz, 2012; Šimunović & Babarović, 2020; VanMeter-Adams et al., 2014). For example, parents who work in STEM fields often provide greater encouragement to their children to go into STEM-related fields (Muenks, Peterson, et al., 2020; Simpkins et al., 2006).

Specifically with respect to research in SEVT, parents’ task values towards STEM fields are positively related to their children developing task values for STEM (Chen, 2001; DeWitt et al., 2013), achievement (Simpson & Oliver, 1990; Sun et al., 2012), and choices to take future STEM courses (Harackiewicz et al., 2012; Rozek et al., 2015; Simpkins et al., 2006; Svoboda et al., 2016). Parents’ expectations and beliefs about their children and their abilities can also shape children’s motivational beliefs (Lee et al., 2020). For example, Parsons and colleagues (1982) found that parents’ beliefs about their students’ mathematics abilities were stronger determinants of their children’s own self-concept and expectancy beliefs than was children’s prior performance in math (Parsons et al., 1982).

One limitation in the literature about parent socialization of motivation is that most research has focused on K-12 education and not college. Existing developmental research suggests that in many life domains, family members may become less influential on students’ attitudes and behavior as students’ progress through adolescence, perhaps due to reduced interaction frequency and dependency (Bokhorst et al., 2010; Larson & Richards, 1991; Spinath & Spinath, 2005). This trend would imply that college students’ beliefs and behaviors are influenced by family members to a lesser extent as they get older. However, the limited work on the role of parents in college illustrates that parents do shape the motivation of college students, such as their beliefs about the malleability of intelligence, at least to some extent (Lin & Muenks, 2024; Starr et al., 2022). It is important to understand how much parents play a role in college students’ motivational socialization, and to explore what this socialization looks like in college, given the crucial socializing role of parents at other ages.

#### Peers & friends

Throughout childhood through emerging adulthood, interactions and relationships with peers, classmates, and friends are widely recognized for their important role on student development and academic achievement (Wentzel, 2005). Positive relationships with peers can promote students’ academic performance (Buhs & Ladd, 2001; Flook et al., 2005; Wentzel, 1998, 2005; Wentzel et al., 2004), act as a source of companionship, help, and emotional support (Parker & Asher, 1993; Riegle-Crumb & Morton, 2017; Wentzel, 1998; Zander et al., 2018), and help develop perspective-taking and empathy, which can foster cooperative, prosocial, and non-aggressive behavior (Card & Little, 2006; Smollar & Youniss, 1989; Wentzel et al., 2007). At the college level, students who are more comfortable with their peers in a group achieve greater content mastery than students who feel uncomfortable with their peers (Theobald et al., 2017). In college chemistry courses, peer learning assistants can provide appraisal and emotional support that increases student engagement in course content (Donis et al., 2024)

With respect to motivational beliefs, relationships with peers and friends are associated with a broad range of value-related outcomes, including liking, positive affect, and enjoyment of school (Ladd et al., 1997; Ryan, 2001). Peers can also impact student’s motivation by acting as role models (Barry & Wentzel, 2006; Schunk & Pajares, 2009), which can influence an individual’s competence-related beliefs (Flook et al., 2005; Robnett & Leaper, 2013), and their sense of belonging, which is associated with their attainment value (Song et al., 2015; Wentzel et al., 2018; Zander et al., 2018). However, peers may not always have a positive influence, especially in college. In some instances, peers can undermine students’ motivation by causing students to make negative social comparisons that threaten competence-related beliefs, devaluing students’ academic achievement or the importance of school tasks, or reducing students’ sense of belonging (Marsh et al., 2008; Urdan & Schoenfelder, 2006). For example, college students who perceive STEM as a cutthroat and highly competitive environment report lower competence related beliefs, lower task values, and higher cost perceptions in STEM (Espinosa, 2011). In undergraduate biology classrooms, students can perceive that male students are the most knowledgeable peers in their class, which may undermine their competence related beliefs and ultimately their career decision making (Grunspan et al., 2016). Peers, then, seem to play a key role in motivational development, but it is not clear whether they are always likely to have a positive influence or the extent to which they may influence students’ career motivation.

#### Mentors, advisors & role models^1^

Supportive relationships with mentors, advisors, and role models can promote individuals’ achievement, success, and motivational beliefs (Crisp & Cruz, 2009; Eby & Robertson, 2020; Gladstone & Cimpian, 2021; Jacobi, 1991; Lawner et al., 2019; National Academies of Sciences, Engineering, and Medicine., 2019; Paglis et al., 2006). Robust meta-analytic evidence shows that mentoring, that is, a relationship between a more experienced individual (i.e., the mentor) aimed at supporting the personal and professional development of a less experienced individual (i.e., the mentee), can lead to a wide range of attitudinal, behavioral, health-related, and motivational outcomes (Eby et al., 2008; Eby et al., 2013). In fact, mentees’ motivation is one of the outcomes most strongly affected by the provision of mentoring support (Blinn-Pike, 2007; Eby et al., 2013). Mentoring may enhance mentees’ motivation by enabling them to acquire new skills and to develop their competence (Kram, 1985; Linnehan, 2003), which in turn, has the potential to beliefs about their competence-related beliefs. At the college level, the provision of mentorship support (e.g., career and psychosocial support) from a post-graduate or faculty mentor can promote students’ career intentions (Aikens et al., 2016; Estrada et al., 2018) and motivation (Hernandez et al., 2017, 2020). Yet, students may also experience negative interactions with their mentors that harm their educational trajectories and career decision making (Authors, 2019; Authors 2021c). Undergraduates who report greater negative experiences with their mentor develop dampened beliefs about the intrinsic and communal value of scientific research, as well as reduced intentions to pursue graduate education and careers in science (Authors, 2024b).

Similar to mentors, role models, or individuals who serve as aspirational examples of behaviors, values, and achievements that others try to emulate, have also been shown to foster students’ motivational beliefs (Gladstone & Cimpian, 2021). For example, Bandura (1977) theorized that role models can positively influence students’ competence-related beliefs via vicarious modeling and verbal persuasion. Role models are also thought to shape students’ expectancies and subjective task values by serving as behavioral models (i.e., demonstrating how to successfully perform a task), representations of the possible (i.e., demonstrating possibilities of success), and acting as a source of inspiration (i.e., demonstrating whether a particular goal is desirable) (Morgenroth et al., 2015). Empirical studies have shown that exposure to role models predicts positively college students’ competence-related beliefs and interest in STEM (Authors, 2024c; Gladstone & Cimpian, 2021; Lawner et al., 2019; Morgenroth et al., 2015). Exposure to role models may be particularly impactful for promoting the motivation of individuals from minoritized and marginalized backgrounds in STEM fields that are both historically and currently dominated by individuals of majority groups (Johnson et al., 2019; Stout et al., 2011).

#### Professors & teaching assistants (TAs)

There is also broad consensus that the quality of students’ relationships with their teachers is pivotal for their academic engagement and achievement (Becker & Luthar, 2002; Hamre & Pianta, 2001; Pianta & Stuhlman, 2004; Skinner & Belmont, 1993; Wentzel, 2009; Wentzel et al., 2010). Teacher behaviors, such as the development and maintenance of supportive relationships (Roorda et al., 2011), transparency in instructional decisions (Murdock et al., 2000; Skinner & Belmont, 1993), providing academic (Newman, 2000) and emotional support (Feldlaufer et al., 1988; Wang & Eccles, 2013), positive student-teacher relationships (Martin & Dowson, 2009), and articulating high expectations while demonstrating interpersonal care through warmth and support (Roorda et al., 2011) have all been shown to predict various aspects of students’ academic motivation (e.g., subjective task value, goal orientation, self-efficacy).

Similar to the influence of family on students’ motivation, the role of teachers, professors, and instructors has been comparatively less studied at the college level compared to the role of teachers in K-12 classrooms. Some emerging research suggests that college-level instructors do continue to play a key socializing role for students. For example, faculty members who are accessible, approachable, respectful, and demonstrate caring attitudes can influence college students’ academic motivation (Hong & Shull, 2010; Komarraju et al., 2010). Additionally, professors’ views on the nature of their student’s ability can have a powerful influence on their students’ performance and sense of belonging in STEM courses (Canning et al., 2019; Hecht et al., 2023; Muenks, Canning, et al., 2020). Finally, research shows that students of instructors who communicate low expectations or messages suggesting that only a few students have the potential to be successful in STEM report lower competence-related beliefs and values for STEM coursework and majors (LaCosse et al., 2021; Muenks, Canning, et al., 2020). It is important to more fully study how college-level instructors might socialize students’ motivational beliefs, especially around career decision-making. In college, it is also important to extend existing work by examining the socializing role of teaching assistants (TAs) alongside instructors. TAs often provide instruction for introductory lecture and/or laboratory courses instead of faculty in STEM fields and often interact with students more frequently than do professors (Rushin et al., 1997; Sundberg et al., 2005).

### Critical Remaining Questions in Understanding How Socializers Shape College STEM Students’ Motivational Development

Few would argue against the significant influence that parents/family members, teachers/professors/TAs, peers/friends, and mentors/advisors/role models can have on students’ motivation. Yet there are several predominant questions that still need to be addressed in the field in order to more fully understand how these socializers shape students’ STEM career motivation and choices.

#### Understanding Socializers in College

One topic that needs more attention is whether the same trends observed for motivational socialization in high school will hold true in the college setting with respect to socialization of career motivation. Much research has examined how, why, and for whom social supports influence an individual’s development during childhood and early/middle stages adolescence (e.g., K-12 schooling), with less research focused on students in college. This is surprising, since and college is a critical developmental and transitional point where students make impactful educational and career related decisions that are influenced by their motivational beliefs (Bubany et al., 2008; Kleine et al., 2021). Without research specifically about socialization of career motivation in college, the field fails to advance our understanding of how and why college students’ make decisions about their careers and how to develop interventions and policy efforts that best support college students making critical career decisions.

#### Understanding Which Socializers are Most Impactful

At the college level, the field knows very little about which social influences are most influential compared to others in shaping students’ motivation and academic decision-making. In K-12 samples, Wentzel’s 1998 seminal study of sixth-grade students’ perceived support from parents, teachers, and peers found that each social influence had a unique role in shaping different types of motivational beliefs (e.g., family cohesion was positively associated with interest and mastery goal orientations and negatively associated with performance goal orientation, but teacher support was positively associated with interest and prosocial goals). In another K-12 study, Song and colleagues (2015) compared the relative contribution of social support provided to 7-9^th^ grade middle school students from parents, peers, and teachers. They reported that while each of the social supports contributed to different motivational outcomes, parental support by far had the broadest and largest influence on students’ motivation compared to the influences of peers and teachers (Song et al., 2015). These findings speak to the critical role of parent influences on motivation for students in K-12 education.

It remains unclear how different types of social supports compare to one another in shaping students’ motivation during college. It is widely believed that the relative importance of parents tends to decrease throughout adolescence, while the relative influence of peers increases (Bokhorst et al., 2010; Larson & Richards, 1991; Spinath & Spinath, 2005). If such trends extend through college that might suggest that parents at some point are no longer the most influential figures shaping students’ motivation. However, given that existing research points to the role of parents continuing to shape motivation in college, it is also plausible that parents continue to be most influential. By understanding which socializers seem to be most influential for college students’ motivational beliefs, such knowledge can advance our understanding of which types of socializers may be the best targets for interventions that can enhance career motivation among college students (e.g., is it better to encourage particular interactions with family members, to provide an intervention via teachers, or to design some kind of peer-based support program?).

#### Understanding How Socializers Shape Motivation

Understanding not only *which* socializers shape college students’ motivation but also *how* they do so is necessary. A given socializer can influence multiple types of motivational beliefs. Yet most existing research at both the K-12 and college level focuses on only one or two motivational pathways at a time (e.g., how does peer support influence competence-related beliefs). As a complement to this approach, it is useful to understand which of several possible motivational beliefs college students most often discuss to be most affected by socializers, and whether socializers shaped those beliefs in a way that attracted them to a particular career or disenchanted them away from a career. This knowledge can help researchers design interventions and/or educational programming that addresses the motivational concerns and beliefs students report to be salient during their social interactions around career decision-making.

In addition to shaping different motivational beliefs, there are multiple different types of socialization behaviors or actions that might lead to changes in the same motivational beliefs. Socialization can take many forms (Cohen & Wills, 1985). For example, emotional support can involve empathic listening, encouragement, and reassurance (i.e., a teacher listening to a student’s concerns about an upcoming exam and reassuring a student). Instrumental support involves socializers providing tangible assistance, advice and guidance, and sharing knowledge and resources (i.e., a mentor or advisor reviewing and providing feedback on an assignment or paper). Social support can also involve a sense of belonging that contributes to social companionship (i.e., strong ties amongst family members). To gain a precise understanding of how socializers shape students’ motivation, it is essential to examine the specific behaviors and supports used by socializers to shape students’ motivational beliefs, alongside understanding which beliefs are affected by social processes.

#### Understanding Which Socializers Shape Motivation for Whom

Educational psychologists have also recognized the need to understand if and how motivational beliefs differ across diverse cultural groups (Strunk & Andrzejewski, 2023). An extensive body of theorical and empirical research indicates that individuals’ cultural backgrounds and identities shape how they make career decisions (Diekman et al., 2010; Emerson & Murphy, 2014; Estrada et al., 2011; Lent & Brown, 2020). To date, however, little research has examined how college students of differing socio-demographic identities (e.g. gender, race, ethnicity, year in college) may uniquely think about how social supports shape their motivation and career decision-making.

Students’ gender, racial/ethnic background, and year in college are all important identities that could lead to unique socialization experiences in STEM career planning. For example, with respect to gender, parents’ beliefs about their children’s ability and career expectations differ based on gender stereotypes, such that parents of males tend to perceive their child as being better at math than the parents of females (Furnham et al., 2002; Yee & Eccles, 1988). These behaviors, in turn, are predictive of students’ actual career decisions years later (Chhin et al., 2008). With respect to race, students of traditionally racially marginalized backgrounds (e.g., American Indian or Alaskan Native, Black or African American, Hispanic or Latino/a) often report interactions with faculty and peers that make them feel intellectually inferior to White students (Carlone & Johnson, 2007; Griffin et al., 2015; Hurtado et al., 2009; Stanton et al., 2022). Asian students, meanwhile, tend to have fewer interactions with faculty members and mentors, and report that their interactions are of lower quality (Aikens et al., 2017; Authors, 2024d; Kim et al., 2009). More broadly, there are multiple structural factors that lead to social environments which can uniquely affect Black students’ perceptions of values (Matthews & Wigfield, 2024). Finally, with regards to year in college, research on the influence of family and peers suggests that as students advance through college, the importance of parents diminishes to some degree, while peer influence takes on a greater role (Bokhorst et al., 2010; Larson & Richards, 1991; Spinath & Spinath, 2005). These relative patterns of importance may affect whom college students perceive as being influential during their career decision making processes, and/or how these socializers are thought to shape motivation.

#### Understanding the Role of Non-Social Supports

Research is also needed to understand how the influence of social supports compares to that of *non-social* supports, or resources that are not derived from social interactions, in shaping college students’ motivation. Non-social supports are also important predictors of student motivation. For instance, internships and job or research experiences during college have a strong impact on students’ motivation (Ceyhan & Tillotson, 2020; Hess et al., 2023; Johari & Bradshaw, 2008; Margaryan et al., 2022; Pender et al., 2010). Students’ experiences in their college and high school courses can similarly shape their interest and competence related beliefs for STEM fields, particularly via the provision of performance feedback (Muenks et al., 2018; Wang & Degol, 2013). Finally, university provided resources, such as career and professional development centers, email listservs, STEM-related clubs or support groups, can also impact college students’ motivation (Chang et al., 2014; Espinosa, 2011). If non-social factors outweigh the role of socializers in shaping students’ motivation, it might suggest that social factors are not the most efficient pathways through which to support career motivational development. Though the primary goal of the present study was to focus on social supports, this study examined social supports in greater depth by comparing their relative influence in shaping career decision-making to the influence of non-social supports.

### The Present Study

In the present study we examined how STEM college students reported that social supports impacted their motivational beliefs regarding their career pursuits. By leveraging quantitative and qualitative data sources and engaging in intensive coding of students’ reports about how social influences shaped their motivation, we hoped to uncover new insights into how different socializers shaped college students’ career motivation in STEM fields. Our specific research questions were:

*Research Question 1:* What social supports (e.g., family, peers, mentors, professors) do STEM college students perceive to be impactful on their career decision-making, both overall and in terms of what factors students report to be most influential?
*Research Question 2:* How do college students’ perceptions of the influence of social supports on their career-decision making differ from their perceptions of the influence of non-social supports (e.g., coursework, career counseling resources, professional development experiences)?
*Research Question 3:* Through what motivational processes do students report that social supports shape their career decision-making (i.e., attraction-related vs. disenchantment related factors; value perceptions vs. competence-related beliefs)?
*Research Question 4:* Are there variations in how these social supports exert their influence on students’ career decision-making depending on students’ gender, race/ethnicity, and year in college?

## Methods

The data in the present manuscript was collected as part of a larger project examining undergraduate students’ perceptions of STEM career decision-making. Other publications have utilized different subsets of this dataset (Authors, under review; Authors, 2024a; Authors, 2025). Each of these studies had different aims, research questions, and utilized completely different subsets of data from the broader dataset compared to the present manuscript (the only overlapping data is on students’ demographics and background characteristics). This study was reviewed and determined to be exempt by the [omitted for blind review] Institutional Review Board ([omitted for blind review].

### Transparency and Openness

We report how we determined our sample size, data exclusions, manipulations, and all measures in the study, and adhered to the journal article reporting standards (Kazak, 2018). Survey data and analysis code are available upon emailed request to the corresponding author. Data was analyzed using R, Version 4.2.2 (R Core Team, 2021). Student text responses are not available due to indirectly identifying information. This study’s design and analysis was not preregistered.

### Context and Participants

We surveyed undergraduate students from one large, public research-intensive university in the Southeastern region of the United States during Spring 2022. To meet the eligibility criteria for the survey, students had to be undergraduate students enrolled in a STEM course. A total of 2,312 students met these criteria and completed the survey. The present study focused exclusively on undergraduates who met the study eligibility criteria and who reported that they intended to major in a STEM field. Of the 2,312 students eligible for the larger study, 83 students were not STEM majors. Excluding these students from the subsequent analyses reduced the final sample size for the present analyses to 2,229 undergraduates. Complete demographic information for undergraduate student participants included in the current study are summarized in Table 1. All participants were treated in accordance with APA ethical guidelines.

**Table 1.** Demographic characteristics of final analytical sample (n = 2,229)

| **Description** | **n (%)** |
| --- | --- |
| **Gender** |  |
| Woman | 1397 (63%) |
| Man | 722 (32%) |
| Non-binary | 24 (1%) |
| Other | 1 (<1%) |
| Prefer not to respond | 72 (3%) |
| **Race/Ethnicity** |  |
| American Indian or Alaskan Native | 8 (<1%) |
| Asian or Asian American | 502 (23%) |
| Black or African American | 164 (7%) |
| Native Hawaiian or Pacific Islander | 11 (<1%) |
| Hispanic or Latino/a | 123 (6%) |
| Middle Eastern | 56 (3%) |
| White or European American | 1466 (66%) |
| Other | 4 (<1%) |
| **Year in College** |  |
| 1<sup>st</sup> Year | 1029 (46%) |
| 2<sup>nd</sup> Year | 490 (22%) |
| 3<sup>rd</sup> Year | 393 (18%) |
| 4<sup>th</sup> Year | 181 (8%) |
| 5<sup>th</sup> Year or beyond | 54 (2%) |
| Other | 2 (<1%) |
| Prefer not to respond | 80 (4%) |
| **College Generation Status** |  |
| No | 1769 (79%) |
| Yes | 391 (18%) |
| Prefer not to respond | 69 (3%) |
| **GPA** |  |
| 3.70 – 4.0 | 1178 (53%) |
| 3.30 – 3.69 | 684 (31%) |
| 3.0 – 3.29 | 188 (8%) |
| < 3.0 | 96 (4%) |
| Prefer not to respond | 83 (4%) |
| **Major** |  |
| Biology | 823 (37%) |
| Computer Science | 293 (13%) |
| Engineering | 371 (17%) |
| Other STEM disciplines (e.g., exercise and sport science, psychology, health promotion, animal science) | 742 (33%) |

*Note.* Students could identify with multiple racial and ethnic categories. Thus, counts will not sum up to 100%.

### Procedure

Students were asked to participate in a survey that focused on their STEM career plans and motivational beliefs. The survey queried students about their satisfaction with their major, long-term career plans, the career navigation resources they utilized, motivational beliefs about career plans, career role models, and background information. Students were recruited through listservs and electronic course announcements from 42 STEM courses across a range of STEM disciplines, representing a total of 26 unique course titles (some courses had multiple sections). We purposefully recruited from courses spanning biology, chemistry, computer science, ecology, engineering, mathematics, microbiology, and physics, to ensure the breadth and diversity of our sample with respect to many different STEM areas. Students in 30 out of the 42 courses received a small amount of extra credit in exchange for their participation in the study. Students on average spent approximately 15-30 minutes completing the survey.

### Measures

#### Survey of overall influences on career navigation

Students were asked to indicate which factors had strongly influenced them in navigating their career decision-making plans (i.e., *“Which (if any) of the following factors have strongly influenced you in navigating your decision-making about your long-term career plans while at [university]? Select all that apply.”)*. They did so by selecting all applicable options from a list generated by the researchers, which consisted of eleven response options. These options were written to reflect a series of factors that may be major influences on students’ career-decision making. This was not at exhaustive list, and we recognize there may be additional factors that could influence students’ career decision making processes, but these factors were selected because they were the ones most often studied in the field.

The social supports on the list included *“interactions with family members,*” “*interactions with classmates or friends,”* “*interactions with course professors or TAs,”* and *“interactions with academic advisors, mentors, or role models you met through [university]”.* The non-social supports on the list were “*courses taken in high school,” “courses taken in college,”* “*research assistantships or research experiences,”* “*job/internship experiences,”* “*university provided resources (e.g., career center),”* “*major provided resources (e.g., email listserv*),” or “*none of the above.”* For parsimony in analyzing the non-social supports and their relative influences, we combined non-social response options in several categories that were conceptually similar together for analyses (coursework in high school and college, research experiences and job/internship experiences), which resulted in a total of nine response categories (four social, four non-social, and one indicating factors not included) that were included in subsequent analyses.

#### Survey of most influential social support on career navigation

After selecting all of the applicable supports that influenced their career decision making, students were asked to select which of these they thought were the most influential support(s) in navigating their career decision-making. Of the factors they originally selected, students could select either one or two factors that they felt were the most influential.

#### Open-ended response regarding how social support affect career navigation

After selecting the most influential factor, students were asked to provide an open-ended response to the question, *“For the factor(s) you just selected, please explain how the factor was influential in shaping your career decision making.”* We examined students’ text responses to identify how these social supports influenced their career decision making (a detailed description of this coding process is described below). Responses ranged from several words to a paragraph of text.

#### Demographics

Students self-reported their gender as man, woman, non-binary, or other^2^. Students also reported their race/ethnicity. Students could identify with multiple racial and ethnic identities, including American Indian or Alaskan Native, Asian or Asian American, Black or African American, Native Hawaiian/Pacific Islander, Hispanic or Latino/a, Middle Eastern, White, or other^2^. Students also reported their year in school (i.e., 1 – 6+ years) and college generation status (i.e., whether or not both of their parents had obtained an undergraduate degree). Students were asked to write in their current majors or intended majors. Student majors were grouped into four broad categories: biology related (i.e., biology, biochemistry and molecular biology, cellular biology, ecology, entomology, genetics, ocean sciences, microbiology, plant biology), computer science related (i.e., computer science, data science), engineering related (i.e., agricultural, biological, chemical, computer systems, electrical/electronics, environmental, mechanical), and other STEM majors (i.e., chemistry, exercise and sport science, psychology, health promotion, animal science). Finally, we collected participants’ self-reported GPA on a 0-4 scale.

### Coding how social supports influenced career decision-making

We conducted qualitative content analyses of open-ended survey responses to characterize how students’ social supports were perceived to influence their career decision making. We analyzed text responses using a three-step coding process involving initial coding, first-cycle coding, and second-cycle coding. First, we began our open coding process (Saldaña, 2013) by reviewing student text responses and making analytic memos noting the similarities, differences, and nuances in how students described the influence of their selected factor(s) on their career decision making. Individual researchers open-coded a subset of student responses, wrote analytic memos describing the processes through which students described the influence of their social supports, and then met as a team to discuss emergent ideas. Through our open coding process, we produced an initial list of ideas that we used to iteratively develop our codebook.

Our initial coding discussions revealed that similar types of motivational processes discussed among student responses in the current study closely mirrored those described in students’ open-ended reflections about career decision-making in previous studies (Authors, 2021a,b). Because of this, we opted to draw from those prior studies’ coding schemes to inform our qualitative coding in the present study. We therefore generated a list of codes for the present analysis using both deductive and provisional coding to refine the a priori set of codes from prior work (Miles et al., 2014; Saldana, 2013).

Our resulting coding scheme based on the a priori research included two components. We first examined whether students perceived that their social supports shaped their career planning in an attraction-related or disenchantment-related way (Table 3). That is, we examined whether students’ social supports were perceived to help them by attracting them towards certain career paths (attraction) or pushing them away from certain career paths (disenchantment). Some students’ responses indicated that their socializers both attracted and disenchanted them from career paths, and such responses were coded as being in both categories.

Second, we relied on the tenets of SEVT (Eccles, 1983; Eccles & Wigfield, 2020) to examine whether students reported that their social supports shaped their career value beliefs or their career competence-related beliefs. Students’ responses were coded as referencing task value beliefs if they referred to themselves or their socializers considering whether a career was useful, interesting, a good fit, meaningful, or otherwise worthwhile. Students’ responses were coded as referencing competence-related beliefs if they referred to themselves or their socializers considering the student’s confidence, capacity, skills, or ability to pursue a certain career or achieve milestones en route to that career (see Table 3 for examples of codes). Students’ responses could fall into both motivational belief categories if applicable.

Three members of our research team applied these provisional coding descriptions to students’ open-ended responses (*n* = 2,059) describing how their most influential social support(s) influenced their decision-making to determine the presence or absence of each category (attraction, disenchantment, task value beliefs, competence-related beliefs). After coding 25 randomly selected responses, the researchers met as a team to discuss, refine, and test the suitability of the codes for describing student responses. The researchers resolved any coding discrepancies by consulting a fourth researcher. This iterative process repeated for three cycles until coding stabilized. Then, three researchers each independently coded a randomly selected subset of 232 student responses (10% of the dataset) to test for reliability, and Cohen’s κs were at an acceptable level for each individual code (κ ranged from 0.77 – 0.91 across codes). Finally, two coders independently coded the rest of students’ responses (*n* coded in this way = 1,827).

We further examined our first cycle codes during a second cycle of analysis. The purpose of our second cycle was to determine the unique processes through which different social supports were perceived as functioning (i.e., which behaviors or interactions socializers used). Our second cycle coding process utilized both pattern coding (i.e., grouping similar data into themes) and axial coding (i.e., determining how similar themes are related and distinct) to identify the dominant ideas students used to describe the unique ways through which social supports shaped their career decision making (Saldaña, 2013; Strauss & Corbin, 1998). As needed, we drew from existing literature to characterize how these supports could act as a source of social influence.

### Analytical Strategy

To address Research Questions 1 and 2, we examined the frequencies of students’ responses regarding the supports they reported as influential in their career decision-making processes. We examined this in terms of which supports students selected most frequently overall, as well as which social supports students selected as being *most influential*.

For Research Question 3, we used our qualitative content analyses of the open-ended responses to characterize how their most influential social supports influenced career decision making. We first examined how social supports influenced students via motivational processes previously discussed in prior literature (i.e., attraction and disenchantment, task values and competence-related beliefs). We then examined what types of specific behaviors and interactions students referenced within each social support as shaping different types of career motivation. We report the frequency of the participants who made comments related to each theme to give a sense of the commonality of the themes within our sample.

To address Research Question 4, we conducted a series of Chi-square tests of independence to determine if there were significant differences in the frequency of selecting a support (overall or as most influential) as a result of students’ sociodemographic characteristics (e.g., gender, race/ethnicity, year in college). Demographic characteristics were coded as follows for the analyses: gender (woman, man); race and ethnicity (Asian, Black or African American, Hispanic/Latine, White); year in college (1-2^nd^ year, 3^rd^ year and beyond). To ensure sufficient statistical power, we omitted non-binary individuals and those who did not disclose their gender from the analyses involving gender. Similarly, we focus analyses on the four racial/ethnic groups with the largest numerical representation in our sample: students from Asian, Black, Hispanic, and White racial and ethnic backgrounds. Given the exploratory nature of this research, we utilized a standard threshold of α = 0.05 and reported all *p* values so that readers can assess these findings independently.

## Results

### What social supports do STEM college students perceive to be impactful on their career decision-making, both overall and in terms of most influential?

Students reported that a variety of supports (*M* = 3.19, *SD* = 1.46) influenced their career decision making overall. Table 2 reports the frequency of how often students selected each of the influences on their decision-making from the researcher-generated checklist. Nearly two-thirds of students selected that interactions with family members played some role in shaping their career decision making (*n* = 1495, or 67.1%). The next most reported influence on decision making was interactions with peers and friends, which was reported by over half of students (*n* = 1195, or 53.6%). Comparatively fewer students reported that interactions with mentors, advisors, and role models (*n* = 740, or 33.2%) and interactions with course professors and TAs (*n* = 577, or 25.9%) had any influence on their career decision making.

**Table 2.** Factors reported by students to be influential in career decision-making.

| **Reason** | **Affected Overall** |  | **Most Influential** |  |
| --- | --- | --- | --- | --- |
|  | *n* | % | *n* | % |
| Courses you've taken in college or high school | 1557 | 69.9 | 734 | 32.9 |
| Interactions with family members | 1495 | 67.1 | 857 | 38.4 |
| Interactions with classmates & friends | 1195 | 53.6 | 362 | 16.2 |
| Professional experiences (job, internship, research experience) | 966 | 43.3 | 554 | 24.8 |
| Interactions with academic mentors, advisors, or role models | 740 | 33.2 | 207 | 9.2 |
| Interactions with course professors or TAs | 577 | 25.9 | 113 | 5.0 |
| University provided resources (career center, email listserv) | 407 | 18.3 | 84 | 3.7 |
| None of the above | 98 | 4.4 | 0 | 0 |

*Note.* Total is $n = 2,229$ . Students could select multiple factors that had an overall influence on their career decision making. Of these factors, they could select up to two factors that were the most influential in affecting their career decision making. Thus, counts will not add up to 100%.

We next examined the social supports that were most frequently selected as being *most influential* in students’ career decision making (Table 2). Again, students mostly commonly indicated that interactions with family members were the most influential in their career decision making (*n* = 857, or 38.4%). 362 students (16.2%) selected interactions with peers and friends as the most influential, 207 students (9.2%) selected interactions with mentors, advisors, and role models as the most influential, and 113 students (5.0%) selected interactions with course professors and TAs as having the greatest influence on their career decision making.

### How impactful are these social supports on students’ career decision-making compared to other non-social supports (e.g., coursework, career counseling resources, professional development experiences)?

In terms of selecting social versus non-social supports (Table 2), over two-thirds of students selected coursework that they had taken in high school or college as influential in their career decision-making (*n* = 1557, or 69.9%). This was a slightly higher frequency than the frequency who selected interactions with family members (67.1%). Nearly half of students reported that professional experiences (i.e., job, internship, research experiences) influenced their career decision making (*n* = 966, or 43.3%). Students selected reasons relating to university provided resources (e.g., career center, email listservs) the least frequently (*n* = 407, or 18.3%). In comparison to social-supports, students selected interactions with peers and friends at a greater frequency than both professional experiences and university provided resources. Interactions with mentors, advisors, and role models, and course professors and TAs were selected less frequently than professional experiences but more often than university provided resources.

In terms of non-social supports selected as *most influential* (Table 2), nearly a third of students indicated that the courses they had taken in high school and college had the largest influence on their decision making (*n* = 734, or 32.9%). This number was smaller than the proportion of students who reported family members to be most influential (38.4%) but larger than the proportion of students reporting other types of social supports to be most influential. A smaller number of students reported that professional experiences were the most influential factors in their career decision making (*n* = 554, or 24.8%). Only 84 students (3.7%) reported university provided resources as being most influential. In summary, students predominantly identified family members as being the most influential in their career decision-making, followed by two non-social factors: coursework and professional experiences. Subsequently, they next most reported three social supports – peers, mentors, and professors – as being most influential. University provided resources were the least frequently selected.

### Through what motivational processes do college students report that social supports shape their career decision-making?

The main themes and example quotations that students referenced regarding how their social supports influence their motivation for career decision-making and frequencies of students reporting different themes, are reported in Tables 3 and 4. Participant quotes have been lightly edited for brevity and clarity.

**Table 3.** Coding scheme and example student text responses.

| **Code** |  | **Definition** | **Sample response** |
| --- | --- | --- | --- |
| Attraction | Task Value-Related Concerns | Influenced students towards a career path by influencing them think about value-related factors | *"I found that I loved to learn and talk about space with my friends. This helped me to come to the realization that space exploration should be something I pursue as a career."* |
|  | Competence-Related Concerns | Influenced students towards a career path by influencing them think about competence-related factors | *"After expressing uncertainty with my original career plan, my family helped me find a career I felt secure and confident in."* |
|  | Other | Influenced students towards a particular career path | *"My dad is a surgeon and I have wanted to pursue a similar career in medicine since I was very young."* |
| Disenchantment | Task Value-Related Concerns | Influenced students away from a career path by influencing them think about value-related factors | *"My family has encouraged me to do what is most interesting [to] me, and I'm realizing that sitting in a lab all day may not be what most interests me."* |
|  | Competence-Related Concerns | Influenced students away from a career path by influencing them think about competence-related factors | *"I was always interested in medicine ever since high school... I didn't think I would have what it takes to get into medical school however, and everyone scared me away from taking hard sciences saying I would fail. So I switched my career plan to business."* |
|  | Other | Influenced students away from a particular career path | *"I currently work at a pharmacy as a pharmacy technician.... I have gotten opinions and advice from people working different positions in this field. After much time, thinking, and taking in all of the information provided, I decided that this field of work wasn't what I wanted to pursue."* |

*Note.* Student responses could be coded as both "other attraction" and "other disenchantment" if their responses contained both elements of these themes. Furthermore, "task value-related concerns" and "competence-related concerns" represent sub-codes of the broader categories of attraction and disenchantment.

**Table 4.** Results of qualitative coding examining how college students’ most influential social support influenced their motivational processes.

| Social support factor | Attraction |  |  |  | Disenchantment |  |  |  |
| --- | --- | --- | --- | --- | --- | --- | --- | --- |
|  | Values | Competence | Other | Total | Values | Competence | Other | Total |
| Family members<br>*n* = 841 | 461<br>(54.8%) | 80<br>(9.6%) | 297<br>(35.3%) | 800<br>(95.1%) | 41<br>(4.9%) | 16<br>(1.9%) | 38<br>(4.5%) | 79<br>(9.4%) |
| Peers & friends<br>*n* = 355 | 181<br>(51.0%) | 37<br>(10.4%) | 124<br>(34.9%) | 323<br>(90.9%) | 27<br>(7.6%) | 14<br>(3.9%) | 26<br>(7.3%) | 53<br>(14.9%) |
| Mentors, advisors, & role models<br>*n* = 202 | 97<br>(48.0%) | 38<br>(18.8%) | 98<br>(48.5%) | 195<br>(96.5%) | 12<br>(5.9%) | 10<br>(4.9%) | 17<br>(8.4%) | 29<br>(14.3%) |
| Professors & TAs<br>*n* = 112 | 62<br>(55.3%) | 5<br>(4.4%) | 44<br>(39.2%) | 109<br>(97.3%) | 5<br>(4.4%) | 4<br>(3.5%) | 4<br>(3.5%) | 12<br>(10.7%) |

*Note.* Total sample for qualitative text analyses *n* = 2,059. The number in parenthesis represents the frequency of the motivational process as reported in student responses for each social support. A small number of students selected a social support as being influential but did not respond to the open-ended question (*n* = 16 for family members, *n* = 7 for peers and friends, *n* = 5 for mentors, advisors, and role models, *n* = 1 for professors and TAs); these participants were excluded from the qualitative analyses. The 'other' category described distinct ways in which a social support shaped attraction or disenchantment that were not related to either task value or competence-related beliefs. The 'total' category indicates those who described at least one reason that the social support that was related to attraction or disenchantment. Percentages do not add to 100% within each group because participants could be classified into more than one category.

#### Family members

Among the students who said family members had the greatest influence on their career decision-making (*n* = 841, or 40.8%), the overwhelming majority described interactions with family members as attracting them towards certain careers (*n* = 800, or 95.1%). Within this group, most students described how their family members helped them think about the positive value afforded by a certain career or how a career aligned with their personal interests (*n* = 461, or 54.8% of students who selected family members as their top influence on career decision-making). Some students recalled that their family members knew them best, which allowed the family members to give advice about what career paths aligned with the students’ interests. Others described how their pursuit of a certain career was perceived as being valuable or interesting through the eyes of their family members. Yet, other students reported that the pursuit of a certain career path would fulfill familial expectations, aspirations, and bring pride to their family. Finally, some students recalled how their family members’ personal experiences influenced their perception of the value in certain career paths. For example, one student said:

> *“Both my mom and best friend suffer from side effects of autoimmune disorders, so I was inspired by them to pursue pharmacy as a career so I can learn for myself what medications have the ability to improve their conditions.”*

Comparatively fewer students reported that their family members helped them realize their competence, ability, or talent for a career path (*n* = 80, or 9.6%). However, some students did articulate that this was a factor in their decision-making. For example, one student described, *“My parents have known me my whole life, so I feel like they have a good perspective and know what I am capable of.”* Other students recalled how their family members believed in their ability which gave them the confidence they needed to pursue a particular career path: *“Many of my family members believe I could be successful with this career.”*

Relative to the number of students who reflected on family influences through a lens of attraction to a new career, a comparatively smaller number referenced family members making them feel disenchanted with particular career paths (*n =* 79, or 9.4% of students who selected family influences as the top factor shaping decision-making). In terms of specific motivational beliefs students discussed within disenchantment, again relatively more students discussed family influences shaping their perceived values (*n* = 51, or 4.9% of those who selected family as their top influence on decision-making), though this was still a small proportion of the overall sample. Students in this category described how their family highlighted the drawbacks of certain careers, describing them as “boring”, “repetitive”, or “unrealistic”, or potentially how pursuing such career paths would make them “unhappy” or that “the stress isn’t worth it.” An even smaller group of students (*n* = 16, or 1.9% of those who selected family as their top influence on decision-making) referenced family members making them feel less competent in a particular career path. Some students described how their family might dissuade them from pursuing a more arduous career due to their own ability or skillset. In other instances, students recalled how their competence was affected vicariously through family members as they saw the challenges family members faced in their own educational and career trajectories. For example, one student explained:

> *“My parents work in the IT industry in various roles, so I’ve heard a lot about their job since young. I used to be a Computer Engineering major as well but I didn’t know if I could handle the workload so I changed to a Computer Science major since I most likely wouldn’t need the hardware knowledge.”*

In terms of specific behaviors or interactions that shaped students’ career motivation, family influences were discussed as being aspirational role models and as giving valuable advice to students, which attracted them to particular careers. For example, one student wrote, “*I grew up farming and working at a cotton gin with my father. His positive influence on my life has led me to pursue a career in the agriculture field.”* A smaller number of students discussed that family members exerted their influence by dissuading or discouraging them from certain career paths, which led to disenchantment.

#### Peers and friends

Relatively fewer students cited peers and friends (hereafter referred to as peers) as the primary influence on their career decision-making (*n* = 355, or 17.2% of the sample) compared to the number of students who selected family members. Similar to family members, students in our sample overwhelmingly reported that their peers exerted their influence by attracting them towards certain careers (*n =* 323, or 90.9% of students who selected this category as their top influence). Just over half of the students who cited the role of peer influence on their career decisions as most influential recalled that interactions with and encouragement from their peers made them see the value of certain careers (*n* = 181, or 51.0%). These students explained that their peers made them aware that certain careers would be more enjoyable than others or that the day-to-day tasks and daily work of some careers would be more closely aligned with their interests. Some students specifically highlighted how their peers influenced them by emphasizing the potential for greater positive emotions and passion in certain career paths. For example, one student stated, *“I needed to choose something that would allow me to be as happy as I could be, and talking to my friends and receiving their feedback has made me want to pursue this if I can.”*

Only a handful of students described that their peers were influential because they helped them feel confident to pursue certain careers (*n* = 37, or 10.4% of those who selected peers as most influential). For instance, some students explained that their peers were influential because they provided emotional encouragement and validation that affirmed they could be successful in their chosen career path. These students also described how their high-performing peers served as role models that they aspired to emulate. By observing their peer’s success or ability, they inferred that they too could be successful in a similar career path. One student explained,

> *“I have friends who are working professionally, and their hard work and intelligence really inspire me. Also, although I am not super far into Computer Science, my perseverance through classes like Calculus 2 and Computer Science, both of which I struggled through but succeeded, make me believe that I can accomplish what I want.”*

A relatively small proportion of students (*n* = 53, or 14.9%) described how interactions with their peers resulted in them being disenchanted with certain careers. Most of the students in this category discussed how their peers influenced their values perceptions (*n* = 27, or 7.6% of the students who selected peers as most influential fell in this category). Some students described viewing their peers’ career goals or current career as being boring, unenjoyable, stress inducing, or unfulfilling which prompted them to question if they were on the right career path. A very small number of students referenced peers disenchanting them away from certain careers by shaping their competence-related beliefs (*n =* 14, or 3.9% of students who selected peers as most influential fell into this category). These were typically described in terms of social comparisons, where students described comparing themselves to their peers and questioning if they had the ability to be successful in a manner similar to their peers. As one student described,

> *“Listening to friends struggling in difficult STEM classes that are required but not necessarily interesting causes me to reflect on whether I actually want to pursue such majors.”*

Family members’ influence in terms of behaviors and interactions was mainly discussed in terms of inspiring future aspirations or offering advice regarding students’ strengths. In contrast, students more often described that peers were influential because they allowed for social comparisons that allowed them to determine if their career goals were attainable or not. Others described how they viewed their peers as role models and aspired to pursue careers that were similar to their peers. Many students also described how their peers provided them with insight on potential careers that they were not previously aware of. For example, one student described,

> *“Talking to peers is more influential because I feel more understood when talking about future careers, worries, and the coursework required for different careers. It is also neat to hear from people my own age that know exactly what they want to do in the future, it is motivating and makes me want to pursue my own career more.”*

#### Mentors, advisors, & role models

Comparatively fewer students referenced mentors, advisors, and role models (hereafter referred to as mentors) as having a profound influence on their career decision-making relative to other social supports (*n* = 202, or 9.8% of the total sample). Similar to how students attributed the role of family members and friends, students also noted that mentors overwhelmingly attracted them towards certain career paths (*n* = 195, or 96.5% of those selecting mentors fell into this category). In terms of specific motivational processes, like with the other social supports, students most often referenced how their mentors attracted them towards certain careers by helping them perceive value (*n* = 97, or 48.0% of students who selected mentors as their top influence fell into this category). Students also expressed how their mentors encouraged them towards career options that aligned with their academic interests or that fit their personality. One student recalled:

*“Because of the passion that I had for video games, one of my advisors advocated that I actually pursue [a career in] it. Before, I was hesitant because of financial reasons.”*

Additionally, some students explained how their mentors affirmed their beliefs and told them to “follow their dreams”, even when these aspirations conflicted with the messages they received from other social supports: *“My advisors have also pushed me to go with the major/career that I am most interested in instead of the career my parents are trying to force onto me.”*

Students in this group also sometimes discussed mentors shaping their competence perceptions when reflecting on attraction-related influences. This occurred more often for mentors relative to how often this was referenced for other social supports (*n* = 38, or 18.8% of students selecting mentors as most influential fell into this category). These students described how mentors played a clarifying role by instilling confidence in their pursuit of careers and making their aspirations feel achievable. Mentors were also describing as providing both technical and emotional guidance and support that made students feel more comfortable with pursuing the training and educational experiences necessary to obtain certain careers. For example, one student wrote,

> *“My advisor was very honest and supportive of me when it came to changing my career. She understood why I was feeling uneasy about my original plan and made me feel confident that I would succeed with my new plan.”*

Like with the other social supports, relatively few students indicated that mentors disenchanted them away from certain careers (*n* = 29, or 14.3%). Of these students, the majority recalled that mentors made them aware of other career paths and advocated that they pursue these careers. In terms of specific motivational beliefs, 12 students (5.9% of the sample of students selecting mentors as most influential) noted how their mentors disenchanted them away from certain careers by influencing their perceived values and interests. Mentors were cited as helping students recognize potential discrepancies between their idealized career aspirations and their personal goals or values. Even fewer students (*n =* 10, or 4.9%) discussed how mentors disenchanted them away from a specific career by influencing their competence-related beliefs. Some of these students described that their mentors made them aware of the demanding and arduous nature of some careers. Others were referenced as bringing attention to various performance issues (e.g., insufficient grades or performance in pre-requisites) that could make it challenging for them to pursue or be successful at their chosen career path. For example, one student recalled:

> *“I used to do fairly well in my courses especially my sciences courses, but I slowly have been having more difficulty grasping the concepts and it has been affecting my grades. I see that that my grades might not make it in such a competitive field, and that is what I have been told by mentors.”*

The precise behavioral processes through which mentors motivated students looked different than it did for family members or friends. Relative to family members or friends, students more frequently spoke about the emotional and psychosocial support offered by mentors that encouraged them to pursue certain career paths. Many students credited their mentors with introducing them to new fields, encouraging them to explore varied opportunities, and offering tangible advice for pursuing certain careers or enhancing their competitiveness for specific career paths. For example, one student explained,

> *“My first mentor introduced to the field of Food Science. She provided me contact information for people within the department and it went from there. My first mentor also helped me to secure my dream internship and told me how she got to where she was, which is exactly where I want to go. Her influence on my career path is incomparable to anything else I’ve experienced.”*

#### Professors & Teaching Assistants

Students reported professors and teaching assistants (hereafter referred to as professors) as having a more modest influence on their career decision making (*n =* 112, or 5.4% of the total sample) compared to other social supports. Professors had an overwhelmingly attraction-focused role in shaping career decision-making (*n* = 109, or 97.3% of students who selected professors as most influential fell into this category). Students often described how their professors broadened their understanding of potentially desirable career options and affirming these decisions in a manner similar to how mentors, advisors, and role models were viewed. For these students, professors often held significant influence as they offered clarity in career decision-making, particular during times when students were grappling with uncertainty about their career plans. One student wrote,

> *“Through talking to different professors and mentors/advisors, I have figured out that there are multiple options as a mechanical engineer. Before speaking to them I honestly did not have a real idea of the possible career options.”*

Over half of the students who attributed the greatest influence on their career paths to professors expressed that these individuals fostered their interest or values in the associated field or line of work (*n* = 62, or 55.3%). Students expressed how they saw their professor as an aspirational role model that was indicative of professionals in their chosen career path or discipline. Other students described ways in which their professors exposed them to career paths that they found interesting, valuable, and in alignment with their identity and values. For example, one student explained,

> *“My favorite class has, without a doubt, been biology with [Professor]… Her class has slowly made me consider career plans outside of the medical field. I feel like so many STEM majors come into college on a default path of pursuing medical school for the prestige, fulfillment of expectations, or money, without really giving it a second thought. I’m trying to break away from that mindset and explore what I truly enjoy.”*

Although somewhat surprising given their role in evaluating students’ performance, students rarely mentioned instances where their professors guided them towards specific careers by affirming their competence- or ability-related beliefs (*n* = 5, or 4.4% of students who selected professors as most influential fell in this category). Students who fell in this category typically referenced how professors who recognized their past performance or accomplishments in their course attracted them towards a career. One student wrote, *“My professors knew from personal experience what my abilities in their class were, and this shaped the advice that they would give me.”*

Of the students who cited their professors as having the largest influence on their career decision-making, only a handful (*n* = 12, or 10.7%) described how professors disenchanted them away from particular career pursuits. Within this group, students described how support from their professors helped them realize they should stop pursuing certain careers. In terms of particular motivational beliefs, very few students (*n* = 5, or 4.4%) indicated that their professors swayed them away from a career path for values or interest related reasons. These students described how their professors highlighted the realities of certain career paths and made them aware of the potential obstacles in career paths that may not be fulfilling or in alignment with their personal interests. One student wrote, *“Talks with my professors in history showed me that I will likely not enjoy a career in academia in general.”* Similarly, only four students (3.5%) described interactions with their professors that disenchanted them via competence-related beliefs. For example, a student recalled how conversations with their professor made them aware of how their current course performance would make it challenging to pursue certain careers.

> *“Many of the courses I am required to take are very difficult and if I do not excel in them, it might negatively impact my chances of getting into veterinary school. Speaking with professors has given me more insight into the profession long term and how adding or choosing other graduate programs to my career plans might align better with my interests.”*

In terms of specific behaviors or interactions that shaped students’ career motivation, professors were described as broadly influential because they shared information that made them more aware of certain careers and inspired them to pursue these career paths. For instance, one student remarked: “*The courses that I have taken in college & the professors I have had, have been extremely inspirational & influential in shaping my career plans.”*

#### Summary of coding results

For all social supports, the overwhelming majority of college students described how their social supports attracted them towards certain career paths. In contrast, only a small number of students described instances where social supports disenchanted them away from certain careers. Additionally, students predominantly described their social supports as influencing their expectancy-value motivational beliefs by shaping their values and interests. Students less frequently described how their social supports influenced their competence-related beliefs. These results demonstrate that students described a variety of social supports as influencing their general motivational processes in similar, if not often overlapping, ways.

Yet, our coding also revealed *distinct* behaviors and characteristics through which each social support seemed to influence students’ career motivation and decision-making. Specifically, family members often exerted their influence by serving as a source of inspiration and providing advice and encouragement as students weighed potential career options. Students also described how family members were uniquely influential because they felt compelled to meet family members’ expectations and goals or pursue certain paths perceived as worthwhile and prestigious by their family. In contrast, students described how they viewed peers as social models whose experiences were highly relevant to their own paths. Through social comparisons peers provided students with means to evaluate their own interests and abilities to succeed in particular career paths. For mentors, students described how they introduced them to previously unknown career paths and provided guidance on how to pursue and effectively obtain their career goals. Mentors also provided emotional support and encouragement when evaluating potential career paths. Finally, professors shared some roles of mentors by introducing students to new career paths and providing guidance about how to pursue certain paths. In addition, students often perceived professors as a source of inspiration or a role model.

### Are there variations in how these social supports exert their influence on students’ career decision-making depending on students’ gender, race/ethnicity, and year in college?

In a final set of exploratory analyses, we examined if students’ thinking about social supports on career decision-making differed as a function of their gender, race/ethnicity, and year in college. In terms of gender differences in social supports (Table 5), both men and women were equally likely to select family members as influential in their career decision-making and to select family as the most influential factor in shaping career decision-making. In fact, both men and women overwhelmingly indicated that family members were the most influential, constituting the highest percentage compared to other supports. However, men were more likely than woman to report that interactions with peers influenced their career decision making, χ^2^ (1) = 5.80, *p* = 0.016, and to indicate that peers were one of the most influential factors to influence their career plans, χ^2^ (1) = 22.15, *p* < 0.0001. In contrast, women were more likely than men to select interactions with mentors as being influential, χ^2^ (1) = 7.00, *p* = 0.008, and to indicate that these interactions were one of the most influential factors to shape their career decision making χ^2^ (1) = 6.23, *p* = 0.012. Women and men were equally likely to indicate that professors affected their career decision-making, both overall and as the primary influences.

**Table 5.** Differences in selection of overall and most influential supports by gender.

| **Affected overall** | **Woman**<br>(*n* = 1397) | | **Man**<br>(*n* = 722) | | $\chi^2$ | *p* |
| --- | --- | --- | --- | --- | --- | --- |
|  | *n* | % | *n* | % |  |  |
| Family members | 947 | 67.8 | 493 | 68.3 | 0.05 | 0.817 |
| Peers or friends | 730 | 52.3 | 417 | 57.8 | 5.80 | **0.016** |
| Course professors or TAs | 371 | 26.6 | 175 | 24.2 | 1.33 | 0.247 |
| Mentors, advisors, or role models | 492 | 35.2 | 213 | 29.5 | 7.00 | **0.008** |
| College or high school courses | 1000 | 71.6 | 489 | 67.7 | 3.38 | 0.065 |
| Professional experiences | 646 | 46.2 | 286 | 39.6 | 8.49 | **0.003** |
| University provided resources | 269 | 19.3 | 121 | 16.8 | 1.97 | 0.159 |
| **Most influential** | **Woman**<br>(*n* = 1397) | | **Man**<br>(*n* = 722) | | $\chi^2$ | *p* |
|  | *n* | % | *n* | % |  |  |
| Family members | 525 | 37.6 | 302 | 41.8 | 3.60 | 0.057 |
| Peers or friends | 192 | 13.7 | 157 | 21.7 | 22.15 | **<0.0001** |
| Course professors or TAs | 67 | 4.8 | 40 | 5.5 | 0.549 | 0.458 |
| Mentors, advisors, or role models | 145 | 10.3 | 51 | 7.1 | 6.23 | **0.012** |
| College or high school courses | 483 | 34.6 | 217 | 30.1 | 4.39 | **0.036** |
| Professional experiences | 343 | 28.1 | 145 | 20.1 | 16.28 | **<0.0001** |
| University provided resources | 52 | 3.7 | 28 | 3.9 | 0.31 | 0.858 |

*Note.* Total *n* = 2119. Values and percentages shown are number of students within a particular group who selected a support. Statistically significant differences ( $p < .05$ ) as indicated by chi-square test are shown in bold font.

For race and ethnicity, there were very few significant differences in the selection of different social supports as being influential (Table 6). Across the four racial and ethnic backgrounds examined, all students reported family members to be highly influential in their career decision making. However, a slightly higher proportion of Asian or Asian American and Black or African American students reported that family members were one of the most influential factors for their career decision making compared to White and Hispanic or Latino/a students, χ^2^ (3) = 13.51, *p* = 0.003. In addition, Asian or Asian American and Hispanic or Latino/a students were more likely to report that peers influenced their career decision making compared to Black or African American and White students (χ^2^ (3) = 11.94, *p* = 0.007), and that peers were one of the most influential factors to influence their career plans, χ^2^ (3) = 10.80, *p* = 0.012. Students across the four racial and ethnic groups were equally likely to select both mentors and professors as affecting and being most influential in their career decision making.

**Table 6.** Differences in selection of overall and most influential supports by race/ethnicity.

| **Affected overall** | **Asian**<br>(*n* = 430) | | **Black**<br>(*n* = 139) | | **Hispanic**<br>(*n* = 67) | | **White**<br>(*n* = 1323) | | $\chi^2$ | *p* |
| --- | --- | --- | --- | --- | --- | --- | --- | --- | --- | --- |
|  | *n* | % | *n* | % | *n* | % | *n* | % |  |  |
| Family members | 308 | 71.6 | 93 | 66.9 | 39 | 58.2 | 893 | 67.5 | 5.78 | 0.122 |
| Peers or friends | 255 | 59.3 | 66 | 47.5 | 44 | 65.7 | 696 | 51.6 | 11.94 | **0.007** |
| Course professors or TAs | 102 | 23.7 | 35 | 25.2 | 19 | 28.4 | 361 | 27.3 | 2.36 | 0.500 |
| Mentors, advisors, or role models | 136 | 31.6 | 47 | 33.8 | 24 | 35.8 | 455 | 34.4 | 1.23 | 0.744 |
| College or high school courses | 279 | 64.9 | 91 | 65.5 | 43 | 64.2 | 963 | 72.8 | 12.71 | **0.005** |
| Professional experiences | 191 | 44.4 | 51 | 36.7 | 23 | 34.3 | 591 | 44.7 | 5.76 | 0.123 |
| University provided resources | 77 | 17.9 | 35 | 25.2 | 19 | 38.4 | 233 | 17.6 | 9.18 | **0.026** |
| **Most influential** | **Asian**<br>(*n* = 430) | | **Black**<br>(*n* = 139) | | **Hispanic**<br>(*n* = 67) | | **White**<br>(*n* = 1323) | | $\chi^2$ | *p* |
|  | *n* | % | *n* | % | *n* | % | *n* | % |  |  |
| Family members | 198 | 46.0 | 60 | 43.2 | 21 | 31.3 | 491 | 37.1 | 13.51 | **0.003** |
| Peers or friends | 92 | 21.4 | 20 | 14.4 | 12 | 17.9 | 196 | 14.8 | 10.80 | **0.012** |
| Course professors or TAs | 17 | 4.0 | 5 | 3.6 | 4 | 6.0 | 70 | 5.3 | 1.93 | 0.586 |
| Mentors, advisors, or role models | 34 | 7.9 | 15 | 10.8 | 9 | 13.4 | 121 | 9.1 | 2.73 | 0.435 |
| College or high school courses | 121 | 28.1 | 39 | 28.1 | 19 | 28.4 | 467 | 35.3 | 9.94 | **0.019** |
| Professional experiences | 98 | 22.8 | 30 | 21.6 | 16 | 23.9 | 353 | 26.7 | 3.84 | 0.278 |
| University provided resources | 16 | 0.70 | 9 | 0.4 | 4 | 0.2 | 47 | 2.2 | 3.70 | 0.295 |

*Note.* Total *n* = 2154. Values and percentages shown are number of students within a particular group who selected a support. Statistically significant differences ( $p < .05$ ) as indicated by chi-square test are shown in bold font.

Finally, the influence of social supports on career decision-making varied as a function of year in college (Table 7). 1^st^ and 2^nd^ year students were significantly more likely to report that family members influenced their career decision making compared to students who were in their 3-6^th^ year of college, χ^2^ (1) = 13.43, *p* = 0.0002, and the same trend held for selecting family members as being most influential in career decision making, χ^2^ (1) = 31.76, *p* < 0.0001. Yet, even with these differences holding true, family members were still reported as being highly influential overall, and as being most influential compared to other supports, across all groups of students. 1^st^ and 2^nd^ years and 3-6^th^ years were equally likely to indicate that friends affected their career decision-making, both overall and as the primary influences. Conversely, a higher proportion of 3-6^th^ year students reported interactions with mentors that influenced their career decision making to some extent, χ^2^ (1) = 5.43, *p* = 0.0197, and that mentors were one of the most influential factors, χ^2^ (1) = 5.44, *p* = 0.0197, than 1-2^nd^ year students. In addition, a higher proportion of 3-6^th^ year students selected interactions with professors as having an influence, χ^2^ (1) = 21.06, *p* < 0.0001, and that this social support was one of the largest influences on their career decision making, χ^2^ (1) = 12.01, *p* = 0.005 compared to 1-2^nd^ year students.

**Table 7.**
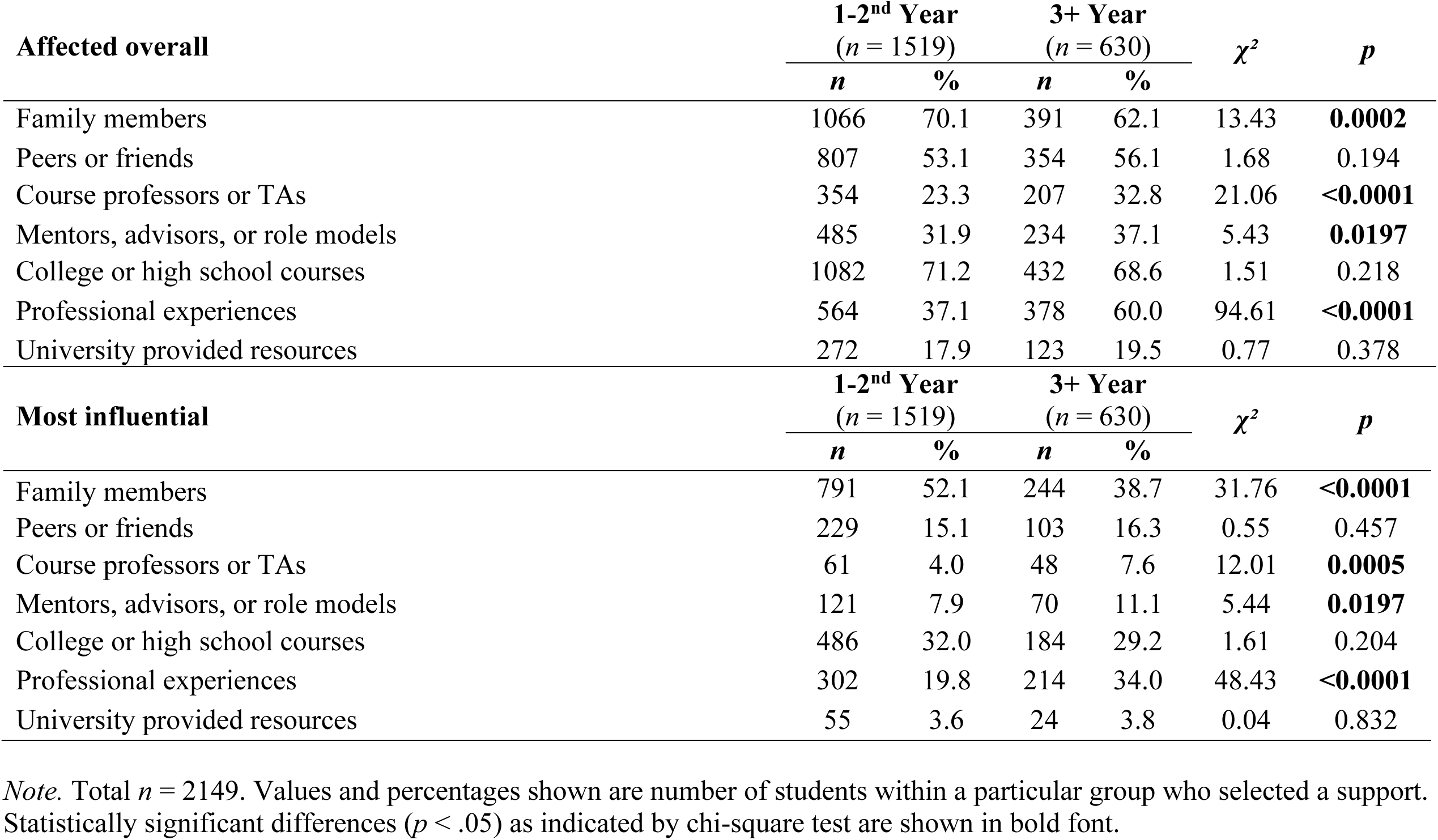
Differences in selection of overall and most influential supports by year in college.

| **Affected overall** | **1-2<sup>nd</sup> Year**<br>(*n* = 1519) | | **3+ Year**<br>(*n* = 630) | | $\chi^2$ | *p* |
| --- | --- | --- | --- | --- | --- | --- |
|  | *n* | % | *n* | % |  |  |
| Family members | 1066 | 70.1 | 391 | 62.1 | 13.43 | **0.0002** |
| Peers or friends | 807 | 53.1 | 354 | 56.1 | 1.68 | 0.194 |
| Course professors or TAs | 354 | 23.3 | 207 | 32.8 | 21.06 | **<0.0001** |
| Mentors, advisors, or role models | 485 | 31.9 | 234 | 37.1 | 5.43 | **0.0197** |
| College or high school courses | 1082 | 71.2 | 432 | 68.6 | 1.51 | 0.218 |
| Professional experiences | 564 | 37.1 | 378 | 60.0 | 94.61 | **<0.0001** |
| University provided resources | 272 | 17.9 | 123 | 19.5 | 0.77 | 0.378 |
| **Most influential** | **1-2<sup>nd</sup> Year**<br>(*n* = 1519) | | **3+ Year**<br>(*n* = 630) | | $\chi^2$ | *p* |
|  | *n* | % | *n* | % |  |  |
| Family members | 791 | 52.1 | 244 | 38.7 | 31.76 | **<0.0001** |
| Peers or friends | 229 | 15.1 | 103 | 16.3 | 0.55 | 0.457 |
| Course professors or TAs | 61 | 4.0 | 48 | 7.6 | 12.01 | **0.0005** |
| Mentors, advisors, or role models | 121 | 7.9 | 70 | 11.1 | 5.44 | **0.0197** |
| College or high school courses | 486 | 32.0 | 184 | 29.2 | 1.61 | 0.204 |
| Professional experiences | 302 | 19.8 | 214 | 34.0 | 48.43 | **<0.0001** |
| University provided resources | 55 | 3.6 | 24 | 3.8 | 0.04 | 0.832 |

*Note.* Total *n* = 2149. Values and percentages shown are number of students within a particular group who selected a support. Statistically significant differences (*p* < .05) as indicated by chi-square test are shown in bold font.

## Discussion

In the present study we examined how and for whom various social supports shape STEM college students’ motivation to make certain career decisions. We draw attention to three major conclusions from our results. First, our findings point to the highly influential role of family members for shaping college students’ career motivation, compared to both other social forms of support and to non-social forms of support, across the college years. Second, results suggest that the most influential social supports students think about for career decision-making are those which attract students towards new careers by helping them think about value-related beliefs. However, within this broad motivational category, different socializers provided different types of behaviors that led to socialization of attraction-related value beliefs. Third, our findings showed that similar social supports were important for various types of students, though some of the specific roles of social supports did differ by students’ year in school and gender. We discuss each of these points further below.

### The importance of family for promoting college students’ career decisions

One major finding from our study is that family members were perceived as one of the most important socializers to affect college students’ career motivational development. We found that the majority of college students in our sample (67.1%) reported interactions with family to play a significant role in their career decisions, making this the second most common factor reported by students, only very slightly less common than coursework taken in college and/or high school (69.9%). Importantly, students also reported family members (38.4%) as the *most influential* factor shaping their career decision-making, a percentage that surpassed the percentage selecting all other influences (both social and non-social). Of critical importance, students reported that family members were most influential among every year in school, gender group, and racial/ethnic group examined. The data clearly demonstrate that family members matter for thinking about how students make career decisions during college.

Prior research underscores the influence of parental support in shaping middle school students’ motivation in comparison to the influences of teachers and peers (Song et al., 2015; Wentzel, 1998). However, support from parents is generally thought to decline during the adolescent years, with support from peers – mainly friends – increasing in importance (Berndt, 1982; Eccles et al., 2013; Larson & Richards, 1991). Our findings add nuance to these points by illustrating that both sets of findings can be true, concurrently, at the college level. That is, parents remain highly influential to students’ career decision-making throughout college, even though students farther along in college more often seemed to report non-parent socializers also were important (see later section of Discussion). More broadly, our findings extend prior research on the relative importance of family members as socializers to the college level and the career decision-making context, a critical point where it is useful to understand how to promote motivation.

These findings also imply that family members may be an under-used resource for helping college students to maintain motivation for STEM career paths or to make informed STEM career decisions. Most educational programs aiming to foster students’ motivation do so by directly intervening with students (e.g., through utility-value interventions or social-belonging interventions; Harackiewicz & Priniski, 2018) or creating instructional environments that support motivation (e.g., through a research experience; Linnenbrink-Garcia et al., 2018). Similarly, most career counseling programs and resources are designed to focus solely on students (Chung & Gfroerer, 2003). Such approaches may unintentionally assume that students are making important career decisions in isolation and/or only within instructional spaces. Our results suggest that it may be worthwhile to design targeted programs or resources to help students interact with their family members in order to gain support and guidance around career development. It may also be important to ensure that family members understand the important role they play in students career decisions and provide family members with tools and resources to provide effective guidance for their children.

Despite family being most important to students, it is important to reiterate that the other social supports examined – peers, mentors/advisors/role models, and professors/teaching assistants – were not *unimportant* in shaping students’ career development. More than half of students reported interactions with peers to be influential in their career decision-making (with about 16% saying they were most influential), and about one-third of students reported mentors, advisors, or role models to be influential (with about 10% saying they were most influential). Though these sources were not the most *frequently* selected in shaping students’ career decision-making in college, these socializers also clearly play key roles in shaping motivation. These findings are important because they suggest that other sources support may complement those received from family. We discuss the specifics of each socializer’s role in more depth below.

### Social supports foster career decisions through attraction and values

We found that students primarily reported their most influential social supports shaped their career decision-making by helping attract them towards certain careers, as opposed to disenchanting them away from the pursuit of certain careers. Similarly, our findings showed that students predominantly described their most influential social supports as shaping their perceptions of task values for particular careers, as opposed to their competence-related beliefs. Notably, these trends appeared across all of the social supports that were discussed. Together, these findings suggest that the strongest social factors influencing STEM career decision-making in individuals’ minds seem to be those which help students see the attractive value of a particular career path.

These findings are novel in that they bridge together two different bodies of empirical literature. In the career decision-making motivation literature, research has illustrated that students often weigh their values and interests as highly salient when making STEM career decisions (Authors, 2021a,b; Harackiewicz et al., 2016; Renninger & Hidi, 2015; Thoman et al., 2014, 2015). Similarly, college students often report both attractive and disenchanting features of new and past careers to shape their career decision-making (Authors, 2021a,b; Authors, 2025; Thoman et al., 2014). However, this work has not focused on how motivational considerations might be socially determined. Similarly, prior work in developmental and educational psychology has shown that social factors play a role in shaping individuals’ motivational beliefs (Song et al., 2015; Wentzel et al., 2010). However, that work has not examined exactly how or through what motivational processes various socializers might shape motivation. By bridging these literatures together, we find that students find it to be especially influential when socializers shape their values and interests, that influential socializers’ roles tend to be much more attraction-related than disenchantment-related in students’ minds, and that these trends seem to be true across a variety of social supports (family members, peers, mentors/advisors/role models, and professors/TAs).

Our results are also consistent with existing social influence theories used to study college student integration into STEM fields, particularly the tripartite integration model of social influence (TIMSI) which examines how individuals orient to a social system (Estrada et al., 2011, 2018). TIMSI posits that three key social influence processes – operationalized as scientific self-efficacy, scientific identify, and science community values - predict students’ integration into the scientific community and ultimately their career interests. Empirical evidence supports the tenets of TIMSI by demonstrating that each of the social influence processes are positively correlated with students’ career interests (Estrada et al., 2011). However, while students’ scientific self-efficacy is important, particularly for performance, it is ultimately students’ scientific identity and values alignment that are uniquely predictive of their behavioral intentions (Estrada et al., 2011; Lee et al., 2024). Our findings are consistent with TIMSI in that we show that influencing agents, such as family, peers, mentors, and professors, have a particularly strong role in shaping students’ values and interests. Thus, these findings suggest that social supports do not exert equal influence on all aspects of students’ motivational beliefs. Rather, they highlight the extent to which motivational processes can be shaped by social supports and the magnitude by which they may be influenced.

Our findings also have important implications for the theoretical model of SEVT, which focuses heavily on social influences. Eccles & Wigfield (2020) recently called for more attention to examine how contextual influences shape students’ achievement motivation. Our results help illustrate exactly how various social supports may relate to students’ motivational beliefs about potential career choices, helping build knowledge of the processes through which achievement motivation develops. Our findings also suggest the novel and intriguing possibility that socializers may have more meaningful roles in shaping students’ value-related beliefs compared to competence-related beliefs. However, this finding would need to be supported by additional studies using varied methodologies and across various contexts before drawing further conclusions.

Results from our study also have direct implications for developing career interventions and resources to aid students in their career decision-making. Our results suggest that career interventions and resources may be especially effective if they leverage the power of socializers to help students think about the attractive, value-related features of particular careers. Extracurricular activities such as career exploration workshops, networking events, and career centers could develop novel materials that highlight how particular careers may align with students’ values and interests and are coupled with support from socializers. For instance, the O*Net Interest Profile (https://www.mynextmove.org/explore/ip) is a valuable assessment tool to help students identify careers that align with their values and interests. While career counselors may use the results to identify careers that align students’ values and interests, encouraging students to discuss these results with family members (or other social supports) may be beneficial for promoting students’ career motivation.

Additionally, novel interventions that integrate existing motivational interventions with social support programs could be enhanced by developing targeted content that emphasizes the specific attractive aspects of careers that are both valuable and interesting to students. Most interventions based in SEVT have been designed to enhance students’ perceptions of utility value and have led to notable effects on student outcomes (Harackiewicz et al., 2016; Hulleman et al., 2010; Hulleman & Harackiewicz, 2009). Furthermore, social support interventions, such as peer mentoring interventions, which pair individuals at similar career stages to provide support, can also be beneficial for career development and planning (Lewis et al., 2017). Connecting utility value interventions around career choices into existing peer mentoring interventions could be a powerful approach for supporting students. Highlighting the attractive and interesting elements of careers to students through existing social programming may improve the efficacy of such programs to more effectively support and motivate students in their career-decision making process.

### Different socializers provide both overlapping and distinct forms of support

Although the motivational influences students discussed (i.e., focusing on attraction, focusing on values) looked similar across socializers, students noted key differences in the specific types of processes and behaviors through which each socializer influenced motivational beliefs. Specifically, family members were commonly described as a source of inspiration and as people who provided relevant advice and guidance to students. Students also described how they felt motivated to adhere to their family member’s perspective and to pursue careers that their family members valued. In comparison, peers were most often described as social models that allowed students to vicariously evaluate their own interests and abilities to be successful in particular career paths. Mentors, advisors, and role models, and professors and TAs, in turn, tended to be described as introducing students to new careers that they were previously unaware of and as sharing guidance and advice on how to pursue these careers. These findings show that different socializers may affect motivation similarly, but they do so by using different types of interactions, behaviors, and experiences. These results are consistent with prior findings suggesting that social supports are not monolithic in their influences on students’ motivational beliefs (Wentzel, 1998). However, they are novel in showing the specific types of social behaviors and interactions that students reported to affect them in the college setting, with respect to career decision-making, and distinctive in using students’ own words to capture how various processes unfolded.

Knowing what types of interactions are likely to be most meaningful in fostering career motivation among different socializers can help inform intervention and future research efforts. That is, rather than assuming a one-size-fits-all approach to social influences, future research and theory can study and intervene on particular types of behaviors for particular socializers, based on which types of behaviors are most often associated with those socializers. Findings also imply that students’ career motivation has the potential to be influenced by multiple socializers via different behaviors and processes, rather than through additive effects of one social process. For example, a student’s decision to pursue a career in a health care field may be influenced by *both* interactions with family members who emphasize the personal importance of the field, while a professor may simultaneously reinforce the students’ sense of competence in the attributes needed for a career in healthcare. Our findings provide insight on which social supports are *most salient* on students’ minds when reflecting on their career decision making, but future research should consider more explicitly how socializers work synergistically during students’ career decision-making.

### Individual differences in social and non-social supports

A final major finding of this study was that students generally reported socializers to have similar effects on motivation regardless of gender, race/ethnicity, or year in school. However, there were some differences that illustrate potentially different pathways of socialization that students might experience as a function of their identities in college. The major difference worth noting is that first and second year students were more likely than students in their third through sixth years of college to select that family members influenced their career decision-making. This is perhaps unsurprising, as by their third year of college, students are more likely to have pursued jobs, internships, and/or coursework that may prompt them to think about their certain career choices more explicitly and critically. Though family members may influence college students at all stages in their learning, the relative role of other supports may become much more important at later stages of education. In terms of other individual differences, we caution readers from overinterpreting these results (see Limitations section) given that our sample of students was somewhat homogeneous in their demographic characteristics (e.g., women, White, in their first year of college). It will be important for future studies to continue exploring how socio-demographics may influence the influence of social supports on students’ motivational beliefs using more robust and rigorous study designs and analytic methods to examine the replicability of these findings.

### Limitations

Our study has limitations and thus provides a starting point for future research. First, data were collected from a single institution with relatively modest representation of students from varied backgrounds and experiences. Thus, our data may not be representative of students at other types of institutions or from differing geographic regions in the United States. Given the evidence that students’ socio-cultural backgrounds can vary widely in the importance they place on social supports, as some may prioritize communal goals or strong family ties, while others might emphasize individual independence, these results may not reflect the full range of ways in which social supports may operate as being influential for students. Our sample size was also limited in ways that precluded our ability to address questions related to intersectionality among the demographic variables. Future research should examine the intersection of students’ gender, racial or ethnic backgrounds, and year in college they relate to their career decision making

Another limitation of this study is that we collected data on some, but not all, forms of social support that may influence college students’ career decisions. In this study, we created a list of possible social and non-social supports theorized to significantly influence students’ career decision-making. However, there may be other important forms of support, both social and non-social, that were not captured in our analysis and may be influential on undergraduates’ career decision-making. Additionally, we examined how students described their social supports as shaping their career decision making in relation to one motivational theory, SEVT. It is likely that various other motivation-adjacent beliefs could be influenced by interactions with social supports and could play a role in influencing students’ career decisions.

Finally, this study provides a snapshot of students’ perceptions of social supports and their roles at a single point in time. This cross-sectional approach does not account for the dynamic nature of these perceptions and how they may evolve longitudinally. While we report how and why students believe that their social support influenced their career decision-making, drawing causal conclusions from qualitative data remains contentious (Maxwell, 2004). We advise readers to be cautious in making causal or directional claims. Finally, the present data are subject to recall bias and the limitations inherent in autobiographical memory, as students attempt to retrospectively make sense of how social supports have shaped their decisions.

### Conclusions

The present study examined how social supports reported by STEM college students impacted their motivational beliefs regarding their career pursuits. Results suggested that family members were the most influential on student’s career decision-making, both overall and as the most influential, relative to interactions with peers/friends, mentors/advisors/role models, and/or professors/TAs. Students predominantly described their social supports as influencing their values and interests, and then to a lesser extent their competence-related beliefs, and these trends were largely robust across all tested demographic groups. The results of this study add to the literature on social influence showing how social supports can influence students career motivation and the processes through which these influences occur.

## Footnotes

1 Mentors, advisors, and role models provide unique support – ranging from guidance, technical support, emotional support, and inspiration – and can act as an example of success, source of guidance and motivation to improve students’ personal and professional development. This distinguishes them from other social supports, such as family or friends, whose roles may be more varied, informal, and less explicitly focused on mentorship or advisory functions. We recognize their distinct functions (Eby et al., 2007), but due to their shared and often overlapping functions, we opted to combine them into one single group to better understand their impact on shaping college students’ motivational beliefs.

2 If students selected “other” when indicating their gender or race and ethnic identities, they were prompted to specify the identity with which they most closely identified in writing.

